# Sex-Specific Links Between Low Choline, Metabolic Dysfunction, and Neuropathology in Obesity: Insights from Humans and the 3xTg-AD Mouse Model of Alzheimer’s disease

**DOI:** 10.1101/2025.02.24.639990

**Authors:** Wendy Winslow, Jessica M. Judd, Savannah Tallino, Lori R. Roust, Eleanna De Filippis, Brooke Brown, Christos Katsanos, Ramon Velazquez

## Abstract

The growing prevalence of obesity, a risk factor for disorders such as Alzheimer’s Disease (AD), raises concerns about the effects on cognitive health. AD currently impacts 6.9 million Americans aged 65 and older and is characterized by the presence of amyloid beta (Aβ) plaques, neurofibrillary tau tangles, and neuroinflammation, all of which contribute to cognitive impairment. Insulin resistance, common in both obesity and AD, disrupts brain glucose metabolism and accelerates neurodegeneration. Understanding the factors that link these conditions could lead to new strategies for combating disease. Notably, the B-like vitamin choline is necessary for fat metabolism and has been shown to help reduce obesity incidence. However, ∼90% of Americans are deficient, and decreases in this nutrient have been associated with cognitive decline. Here, we examined circulating choline levels, inflammation, and metabolic dysfunction in human participants with obesity (BMI > 30) compared to normal BMIs (18.5–24.9), as well as in 3xTg-AD mice, an AD model, fed a choline-deficient diet throughout adulthood. Our results revealed that obese participants exhibited significantly lower circulating choline levels compared to those with a healthy BMI. Lower choline levels correlated with higher %Body fat and increased markers of insulin resistance. Elevated inflammatory cytokines in obese participants were also seen in 3xTg-AD mice on a choline-deficient diet, which exhibited significant weight gain and metabolic dysfunction. AD-like pathology was also exacerbated in choline deficient 3xTg-AD mice. These findings underscore the relationship between low choline levels, obesity, insulin resistance, and cognitive decline risk. Adequate choline intake may mitigate the risk of obesity, potentially preventing cognitive decline and associated diseases.

**Highlights:**

- Obesity is linked to increased insulin resistance (IR) and systemic inflammation, both of which are recognized risk factors for Alzheimer’s disease (AD).
- Women exhibit lower circulating choline levels compared to men, and obese individuals display significantly lower choline levels than those with a healthy BMI.
- Lower circulating choline levels are linked to a higher body fat percentage, increased markers of IR and liver dysfunction, as well as heightened systemic inflammation.
- 3xTg-AD mice on a choline-deficient diet experience considerable weight gain, metabolic dysfunction, heightened systemic inflammation, and AD-like pathology, resembling the conditions observed in obese human participants.

## Introduction

Obesity is a global health pandemic, and its prevalence is on the rise [1]. In the United States (U.S.), from 1999-2018, obesity increased from 30.5% to 42.4% [1]. Notably, obesity is higher in women than men, and this discrepancy increases with age [1]. Obesity is defined as a body mass index (BMI) of 30.0 kg/m² or higher and is associated with a range of chronic health issues, including increased risk of type 2 diabetes (T2D), chronic systemic inflammation, and neurodegenerative disorders [2]. Gaining a deeper understanding of how obesity contributes to these conditions and shared underlying mechanisms is crucial, as early intervention could help prevent or reduce the risk of developing more severe and debilitating sequelae.

Among the metabolic dysfunctions that can accompany obesity, insulin resistance (IR) is one of the most insidious. Obesity contributes to IR by increasing visceral fat which secretes pro-inflammatory cytokines, disrupting normal metabolism [3]. Insulin is an essential hormone secreted by β-cells of the pancreas [4], playing a key role in promoting glucose uptake into cells, especially muscle and adipose tissues. It does so by binding to insulin receptors on the cell surface, which activates a series of intracellular signaling pathways, most notable the PI3K-AKT pathway [4,5]. Insulin maintains glucose homeostasis by inhibiting hepatic glucose production and promoting glycogen, lipid, and protein synthesis [5]. However, IR occurs when cells in the body become less responsive to insulin [3]. Obesity, poor diet, physical inactivity, and genetic predisposition increase the likelihood of IR development [3,5,6]. IR is associated with lipid accumulation in the liver and skeletal muscle, endoplasmic reticulum stress, and chronic inflammation, resulting in hyperglycemia, dyslipidemia, and hypertension, which collectively contribute to the development of metabolic syndrome [5]. Gaining a deeper understanding of the causes of IR could provide insights into preventive strategies to mitigate its adverse health impacts.

IR is a major risk factor for Alzheimer’s disease (AD), which is the most common form of dementia, and which currently affects 6.9 million adults aged 65 and older in the U.S. [7]. AD is marked by the buildup of amyloid beta (Aβ) plaques, neurofibrillary tau tangles, and chronic systemic inflammation, all of which play a role in the gradual loss of memory and other cognitive abilities [7]. Interestingly, the connection between AD and IR is so strong that AD is frequently referred to as a neuroendocrine disorder and dubbed “type 3 diabetes” [4,6]. Brain insulin plays a critical role in regulating appetite, cognitive function, and overall energy balance through its effects on the central nervous system, with less direct involvement in glucose uptake within the brain itself compared to its effects peripherally; brain IR contributes to obesity by disrupting the brain’s ability to regulate appetite and energy expenditure [8]. Furthermore, impaired insulin signaling contributes to cognitive decline by promoting Aβ plaque accumulation, tau hyperphosphorylation, and ultimately neurodegeneration—all hallmarks of AD [8,9]. In the brain, the PI3K/AKT/GSK-3β signaling pathway regulates tau phosphorylation, neuronal survival, and Aβ clearance [10], a pathway that is dysregulated by IR. Therefore, IR – often induced by obesity – facilitates the progression of AD pathological features, highlighting IR as a key target for interventions designed to prevent the development of AD.

Since lifestyle factors – such as diet – can contribute to obesity, dietary intake of the nutrient choline is of interest as it can mitigate IR, inflammation, and AD pathology [11–21]. Choline is an essential nutrient that plays key roles across a wide range of organ systems, including in fat metabolism, serving as a precursor of acetylcholine, and in building cell membranes [12,22]. While 30% of the required choline is produced endogenously by the enzyme phosphatidylethanolamine N-methyltransferase (PEMT), the remaining amount must be obtained from diet [22]. Alarmingly, 90% of Americans fail to meet the recommended daily intake of choline, which can compromise a wide range of metabolic functions [23]. Converging evidence from human and rodent studies highlights the association between higher dietary choline intake and a lower risk of developing IR [13] and T2D [15]. Choline intake in healthy adults is also associated with reduced inflammation. For example, those consuming more than 310 mg/day show 26% lower IL-6 and 6% lower TNF-α levels, both of which are pro-inflammatory cytokines related to metabolic dysfunction [17]. Similarly, higher levels of circulating choline are associated with lower inflammation, including reduced TNF-α, in patients with either mild cognitive impairment or AD [11]. 3xTg-AD mice fed a choline deficient diet exhibit higher levels of TNF-α [11], while dietary choline supplementation reduces inflammation and AD-like pathology in APP/PS1 mice [18]. This underscores the intricate connection between choline deficiency and obesity, inflammation, and neurodegenerative disorders such as AD, highlighting that preserving choline status may help mitigate the interconnected issues of IR, inflammation, and AD.

The aim of this study is to explore the relationship between early to mid-life obesity, metabolic dysfunction, circulating choline levels, and inflammation profiles in blood plasma from individuals with obesity (BMI > 30 kg/m²) compared to age-matched individuals with healthy BMIs (18.5-24.9 kg/m²). Furthermore, we examined choline levels, glucose metabolism, and inflammatory cytokine profiles in a mouse model of AD on a choline-deficient diet, enabling us to interrogate parallels between obese individuals without a clinical AD diagnosis and an AD model on a choline-deficient diet. We hypothesized that obese participants would exhibit low circulating choline levels, which would correlate with increased IR, systemic cytokine levels, and proteins associated with liver dysfunction, consistent with the changes observed in choline-deficient 3xTg-AD mice.

## Materials and Methods

### Human Participants

Subjects were recruited through online and paper advertisements from the greater Phoenix metropolitan area in Arizona and assigned to the appropriate group based on their body mass index (BMI). Participants included individuals with BMI < 24.9 kg/m^2^ (Healthy) and individuals with BMI > 30 kg/m^2^ (Obese). The human studies were performed after obtaining approval from the Institutional Review Board at Mayo Clinic (IRB #: 20-003294). Written consent was obtained from each participant prior to any study procedures. Study participants were determined to be healthy based on medical history, routine physical examination, electrocardiogram, standard blood tests, and urinalysis. Those with diabetes, history of liver, renal, or heart disease were excluded from the study, as well as those that smoked, participated in a weight-loss regimen, took nutritional supplements, or used prescription or over-the-counter medications. Body composition, including percent body fat, was measured using bioelectrical impedance analysis (BIA) (BIA 310e, Biodynamics Corp., Shoreline, Washington). Blood samples were collected from participants after a 12-hour fasting period and processed at the Mayo Clinic Laboratory, including for measurements of A1C and triglycerides.

### Animals

Mice were group-housed (4-5 per cage) and kept on a 12-h light/dark cycle at 23°C with *ad libitum* access to food and water. All animal procedures were approved in advance by the Institutional Animal Care and Use Committee of Arizona State University. 3xTg-AD mice express a Swedish mutation of the amyloid precursor protein (APP), a mutated presenilin 1 (PS1 M146V) to accelerate amyloid deposition, and a mutated human tau (P301L), leading to tau pathology [11,12,24]. C57BL6/129S6 mice were used as non-transgenic (NonTg) controls. In line with previous research showing inconsistent pathology in male 3xTg-AD mice, only female mice were used in this study. [11,12,24]. 3xTg-AD mice develop Aβ starting at 6 months and pathological tau by 12 months [11,12,24]. Genetic drift is seen in the 3xTg-AD model and our lab backcrosses our mice as recommended by the Jackson Laboratory to minimize such effects. 3xTg-AD and NonTg mice were randomly assigned to one of two diets at 3 months of age; a standard laboratory AIN76A diet (Envigo Teklad Diets, Madison WI) with adequate choline (ChN; 2.0 g/kg choline chloride; #TD.180228), or an AIN76A choline-deficient (Ch-; 0.0 g/kg; #TD.110617) diet, as previously described [12]. The four mouse groups were NonTg ChN, NonTg Ch-, 3xTg-AD ChN, and 3xTg-AD Ch- (*n* = 4/group). Where noted, mice used for analysis were a subset of the previously published data in Dave et. al. 2023 [12]. Choline levels, inflammatory panels, and pathology of this specific subset have not been previously reported.

### 3xTg-AD mouse weight and glucose tolerance (GTT) test

Weight was measured biweekly. A GTT was performed, as previously described, prior to euthanasia [12,24]. Briefly, animals were fasted overnight for 16 h, and baseline fasting glucose levels were taken. Animals then received a 2.0 g/kg glucose intraperitoneal (i.p.) injection and blood glucose was sampled from the tail using a TRUEtrack glucose meter and TRUEtrack test strips (Trividia Health) at 15, 30, 45, 60, 90, 120, and 150 min following the injection. The data used in this analysis were a subset of mice in Dave et al., 2023 [12].

### Blood and tissue collection

At 12 months of age, 3xTg-AD and NonTg mice were fasted for 16 h, and 150-200 μL (≤ 1% of the subject’s body weight) of blood was collected via the submandibular vein and placed into EDTA-lined tubes (BD K_2_EDTA #365,974). Tubes were inverted eight times to assure anticoagulation, kept on ice for 60-90 min, and then centrifuged at 455x g for 30 min at 4 °C to separate phases. The top layer was collected and frozen at −80 °C prior to use. Three to 14 days later, mice were sacrificed via CO_2_ inhalation and brains were harvested. The cortex was dissected out from one hemisphere, flash-frozen with dry ice, and stored at −80°C. Extracted tissue was homogenized in a T-PER tissue protein extraction reagent supplemented with protease (Roche Applied Science, IN, United States) and phosphatase inhibitors (Millipore, MA, USA). The homogenized tissues were centrifuged at 4°C for 30 min, and the supernatant (soluble fraction) was stored at −80°C. The remaining pellet was further homogenized in 70% formic acid and then centrifuged at 4°C for 30 min. The supernatant (insoluble fraction) was collected, neutralized, and stored at −80°C.

### Multiplex Cytokine analysis

To assess the levels of cytokines in humans and mice, we used a Bioplex human cytokine 11-plex kit (Bio-Rad, cat. no. #12003080) and a Bioplex mouse cytokine 23-plex kit (Bio-Rad cat. No. # M60009RDPD) as previously described [25]. Briefly, plasma samples were diluted 1:10 in sample diluent, added in duplicate to a 96-well plate, and assayed according to the manufacturer’s instructions. Measurements and data analysis were performed using the Bio-Plex Manager software (version 6.1). Standard curves were plotted using five-parameter logistic regression and concentrations were calculated accordingly. Cytokine levels that were not detected based on manufacture limits were not included in the analysis.

### ELISA and Plate Assays

Plasma glucose concentration was measured using an automated glucose analyzer (YSI 2300, Yellow Springs, OH). Plasma insulin was measured using a commercial ELISA assay (Alpco Diagnostics Cat# 80-INSHU-E01.1) and used to calculate the Homeostatic Model Assessment for IR (HOMA-IR). Choline levels were measured with a commercially available colorimetric kit (Abcam, ab219944), following manufacturer guidelines, as previously described [11,12]. Aldolase B activity was also assessed using a colorimetric assay kit from Millipore Sigma (MAK223). Commercially available ELISA kits were used to measure sorbitol dehydrogenase levels in human plasma samples (Abcam ab233613), following the manufacturer’s guidelines. Amyloidosis and hyperphosphorylated tau markers were measured using ELISA kits (Invitrogen-ThermoFisher Scientific) to quantify cortical levels of soluble and insoluble Aβ40 (Cat# KHB3481), Aβ42 (Cat# KHB3441), and phosphorylated Tau (pTau) at T181 (Cat# KHO0631) and S396 (Cat# KHB7031), as previously described [11,12,25].

### Statistical Analysis

Data was analyzed using GraphPad Prism (version 10.3.1). Group differences for mouse pathology were analyzed using unpaired t-tests. All other group differences were analyzed using two-way factorial ANOVAs (BMI classification by sex in humans, genotype by diet in mice) followed by recommended post-hoc comparisons when appropriate. Assumptions of normality were met. Linear correlations were calculated using Pearson’s r coefficient. Significance was set at *p* < 0.05.

## Results

### Obese participants exhibit metabolic disturbances, including IR

Subject profiles are detailed in Table 1. Participants were age-matched and balanced for sex (Healthy Males *n* = 7, *M*_age_ = 35.86 years; Healthy Females *n* = 8, *M*_age_ = 34.13 years; Obese Males *n* = 8, *M*_age_ = 35.14 years; Obese Females *n* = 7, *M*_age_ = 29.14 years). Obese participants had significantly higher body weight (*F*_(1,26)_ = 52.54, *p* < 0.0001) and BMI (*F*_(1,26)_ = 141.6, *p* < 0.0001) than healthy participants. Obese participants exhibited higher %Body Fat (*F*_(1,26)_ = 76.11, *p* < 0.0001) than healthy participants, and women had higher %Body Fat (*F*_(1,26)_ = 83.78, *p* < 0.0001) than men. There were also notable differences in common markers of metabolic health. Triglyceride levels were higher in obese (*F*_(1, 25)_ = 4.688, *p* = 0.0401) than in healthy participants, and an interaction between BMI classification and sex (*F*_(1,25)_ = 5.826, *p* = 0.0234) showed that obese men had higher triglyceride levels than both healthy men (*p* = 0.0028) and obese women (*p* = 0.0063). Fasting glucose levels and A1C – a metric of average glucose levels over the last three months [26] – were higher in obese participants but did not reach statistical significance. Insulin levels (*F*_(1, 26)_ = 12.72, *p* = 0.0014) and HOMA-IR (*F*_(1, 26)_ = 13.37, *p* = 0.0011) – a metric of IR [27] – were both higher in obese participants compared to healthy participants, indicating IR in obese participants. These results highlight metabolic disturbances in obese participants.

**Table 1.**
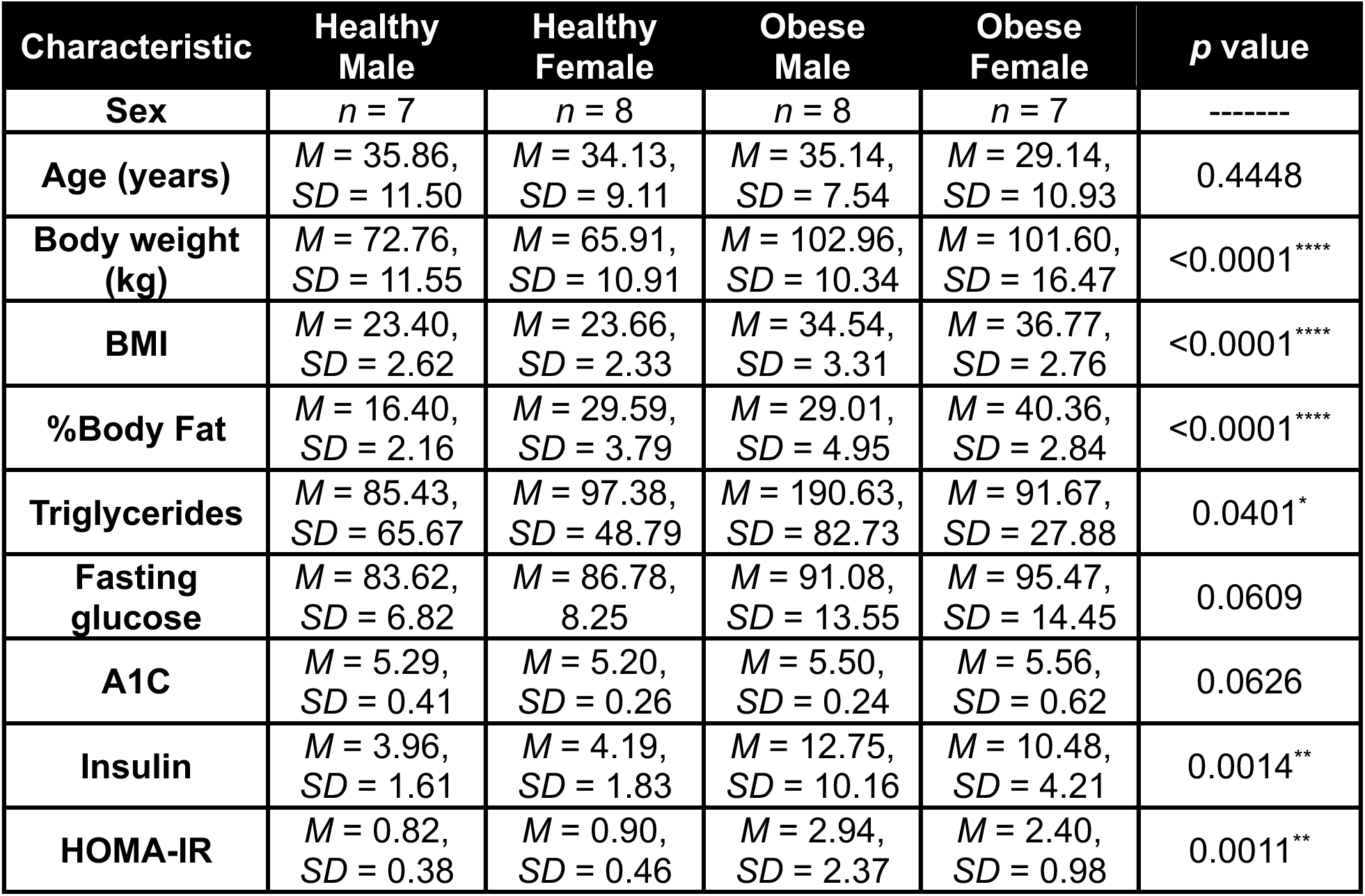
Characteristics of participants classified as controls with a healthy BMI compared to those classified as obese BMI. Blood measures reported were taken after a 12-hour fasting period. *Abbreviations:* Mean (*M*), Standard deviation (*SD*), Body mass index (BMI), Homeostatic Model Assessment for Insulin Resistance (HOMA-IR).

### Plasma choline is reduced in obese participants and correlates with markers of metabolic dysfunction

We next measured blood plasma choline levels and found that obese participants had lower choline levels than healthy participants (Fig. 1A; *F*_(1,25)_ = 102.4, *p* < 0.0001). Notably, we found lower choline levels in women than in men (*F*_(1, 25)_ = 6.310, *p* = 0.0188), consistent with reports highlighting that women consume less dietary choline than men [23]. Next, we ran correlations between choline levels and metabolic metrics. Higher BMI (*r*_(28)_ = −0.8293, *p* < 0.0001) and %Body Fat (*r*_(28)_ = −0.6735, *p* < 0.0001) were correlated with lower plasma choline (Fig. 1B, C). There were no significant correlations between choline levels and glucose or A1C (Fig. 1D, E). Insulin (Fig. 1F; *r*_(28)_ = −0.4720, *p* = 0.0097) and HOMA-IR (Fig. 1G; *r*_(28)_ = −0.4092, *p* = 0.0275) were negatively correlated with choline levels, highlighting that as choline levels go down, these metabolic measures increase. Together, these results demonstrate that obesity and IR metrics correspond with lower circulating choline levels, which may be tied to dietary intake of choline.

**Figure 1.**
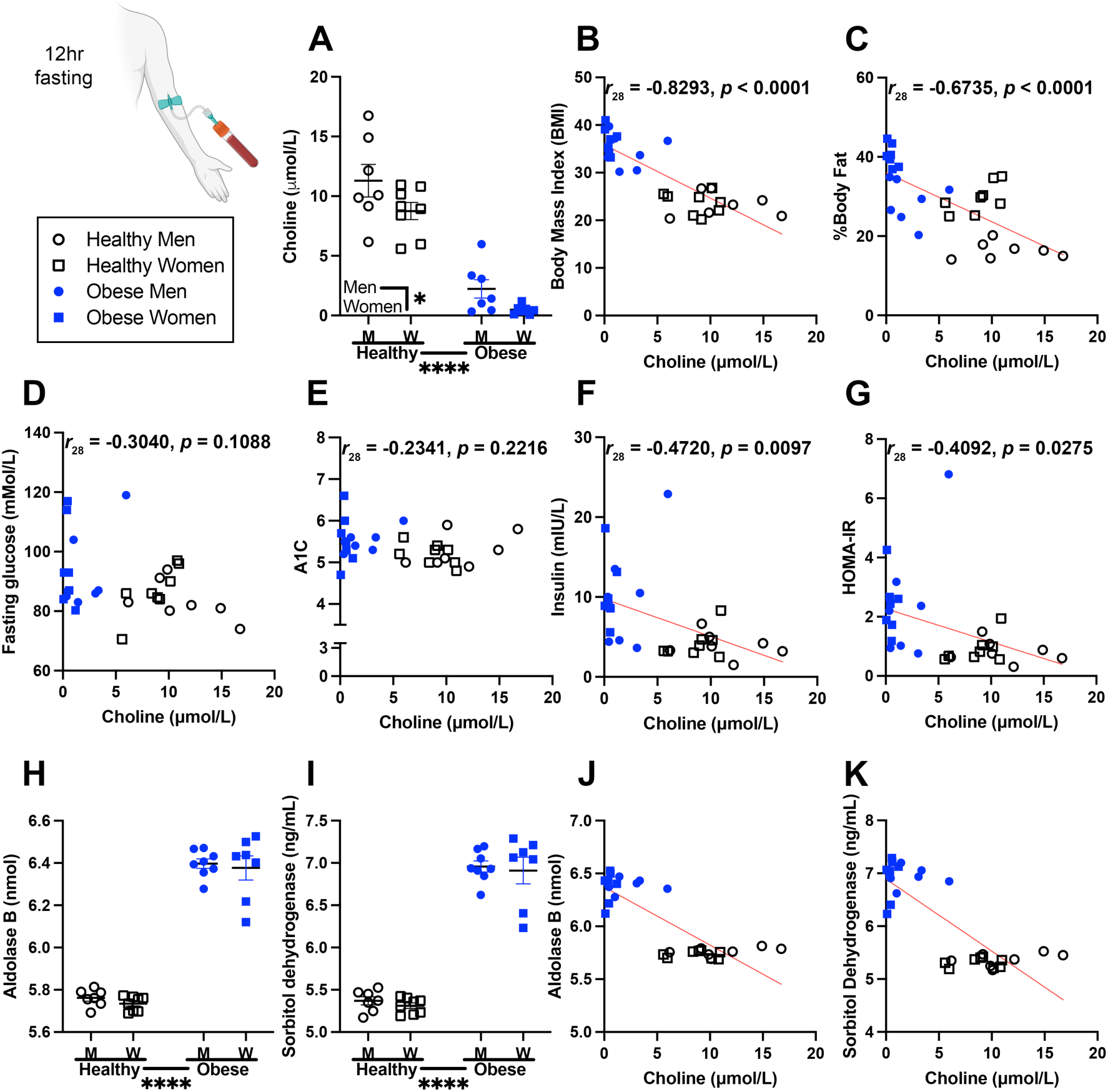
Plasma choline was reduced in obese participants and negatively correlated with multiple metabolic metrics. **(A)** Plasma choline levels were reduced in obese participants compared to healthy weight participants and were lower in women than in men. **(B)** Body mass index (BMI) and **(C)** %Body Fat were negatively correlated with plasma choline levels. **(D)** Fasting glucose and **(E)** A1C were not significantly correlated with plasma choline levels. **(F)** Insulin and **(G)** HOMA-IR were negatively correlated with plasma choline. Liver enzymes are elevated in obese participants compared to healthy participants. **(H)** Aldolase B and **(I)** sorbitol dehydrogenase levels were elevated in obese participants compared to healthy participants. Elevated **(J)** aldolase B and **(K)** sorbitol dehydrogenase levels correlated with lower choline levels. Data are reported as means ± SEM. \**p* < 0.05, **P < 0.01, \*\*\**p*<0.001, \*\*\*\**p* < 0.0001.

### Enzymes that are indicative of liver dysfunction are elevated in the plasma of obese compared to healthy participants

We recently observed that enzymes aldolase B and sorbitol dehydrogenase are significantly elevated by a choline deficient diet in both NonTg (Log2 Fold Change = 2.25 for aldolase B; 1.62 for sorbitol dehydrogenase) and 3xTg-AD (Log2 Fold Change = 1.65 for aldolase B; 1.67 for sorbitol dehydrogenase) mice [12]. Aldolase B and sorbitol dehydrogenase are enzymes involved in carbohydrate metabolism and elevations in these enzymes in blood are indicative of liver injury and dysfunction [28–30]. Aldolase B is expressed in the liver and contributes to the metabolism of fructose [30]. We found elevations in obese participants compared to healthy-weight participants (Fig. 1H; *F*_(1,26)_ = 434.4, *p* < 0.0001). Similarly, sorbitol dehydrogenase, which participates in the conversion of glucose into fructose [30], was also elevated in obese participants relative to healthy weight participants (Fig. 1I; *F*_(1,26)_ = 344.5, *p* < 0.0001). Moreover, aldolase B (Fig. 1J; *r*_(28)_ = −0.8275, *p* < 0.0001) and sorbitol dehydrogenase (Fig. 1K; *r*_(28)_ = −0.8182, *p* < 0.0001) were both negatively correlated with plasma choline. Together, this indicates IR and liver dysfunction in obese participants compared to healthy participants, and, since choline is important for maintaining liver health, suggests that low circulating choline levels may contribute to this dysfunction.

### Pro-and anti-inflammatory cytokine levels are elevated in obese participants

We next measured eleven cytokines that are known to be associated with IR and AD [3,11,25,31–34] and found that all were significantly elevated in obese participants (Fig. 2A). The inflammatory cytokines IFN-γ (Fig. 2B; *F*_(1,26)_ = 48.78, *p* < 0.0001), IL-1β (Fig. 2C; *F*_(1, 26)_ = 94.23, *p* < 0.0001), and IL-2, (Fig. 2D; *F*_(1, 25)_ = 64.00, *p* < 0.0001) were significantly elevated in obese participants. Additionally, there was a trend of elevated IL-1β in women (*F*_(1, 26)_ = 4.081, *p* = 0.0538). Anti-inflammatory cytokines IL-4 (Fig. 2E; *F*_(1, 25)_ = 45.44, *p* < 0.0001) and IL-5 (Fig. 2F; *F*_(1, 25)_ = 85.34, *p* < 0.0001) were both significantly elevated in obese participants, indicating a potential compensatory mechanism attempting to restore homeostasis [3]. IR promoting cytokines IL-6 (Fig. 2G; *F*_(1, 26)_ = 84.29, *p* < 0.0001) and IL-8 (Fig. 2H; *F*_(1, 26)_ = 61.90, *p* < 0.0001) were significantly elevated in obese participants. IL-10, which, depending on the target tissue, can increase or decrease insulin sensitivity, (Fig. 2I; *F*_(1,26)_ = 93.83, *p* < 0.0001) and IL-12 p70, the active form of IL-12, (Fig. 2J; *F*_(1, 24)_ = 44.31, *p* < 0.0001) were significantly elevated in obese participants. The anti-inflammatory cytokine IL-13 (Fig. 2K; *F*_(1, 24)_ = 44.27, *p* < 0.0001) was also significantly elevated in obese participants. Lastly, inflammatory cytokine TNF-α (Fig. 2L; *F*_(1, 26)_ = 70.30, *p* < 0.0001) was significantly elevated in obese participants. Together, these results highlight heightened inflammation in obese participants who also display IR and low circulating choline levels.

**Figure 2.**
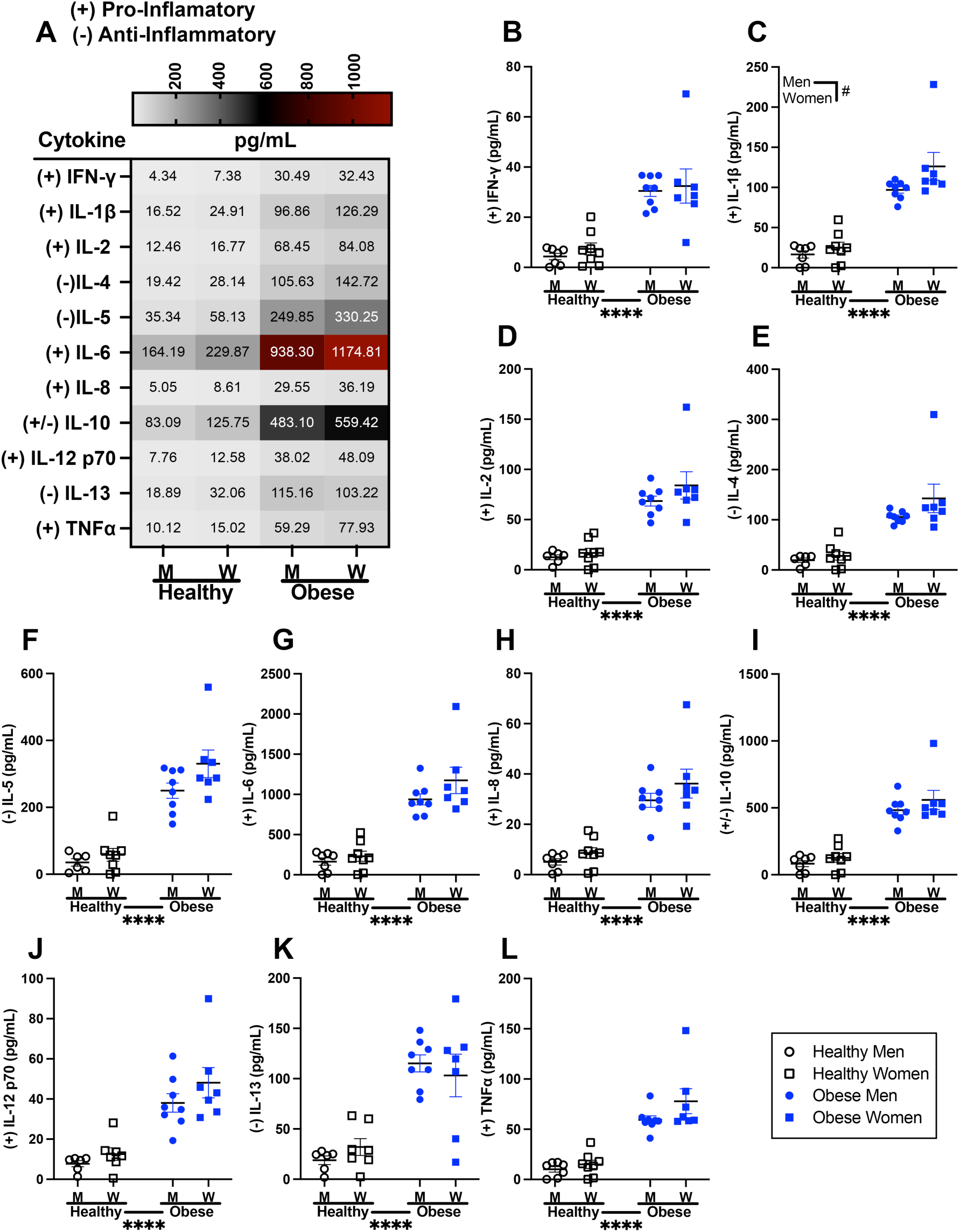
Eleven cytokines were elevated in obese compared to healthy participants. **(A)** A heat map of the cytokine panel illustrating the cytokines that were altered in obese participants. **(B)** IFN-γ, **(C)** IL-1β, **(D)** IL-2, **(E)** IL-4, **(F)** IL-5, **(G)** IL-6, **(H)** IL-8, **(I)** IL-10, **(J)** IL-12 p70, **(K)** IL-13, and **(L)** TNF-α were higher in obese participants than in healthy weight participants. Data are reported as means ± SEM. \**p* < 0.05, \*\**p* < 0.01, \*\*\**p*<0.001, \*\*\*\**p* < 0.0001.

### A choline-deficient diet in mice induces obesity and glucose metabolism impairments that parallel observed changes in human obese participants

At 3 months of age, 3xTg-AD and NonTg mice were assigned to either a choline-sufficient diet (ChN) or a choline-deficient diet (Ch-) and maintained on these diets until 12 months of age (n = 4/group; Fig. 3A). We found that body weight increased over the duration of the study (Fig. 3B). Notably, at 7 months of age, we found significant main effects of genotype (*F*_(1, 12)_ = 10.67, *p* = 0.0068) and diet (*F*_(1, 12)_ = 23.14, *p* = 0.0004), where % change in body weight was higher in the 3xTg-AD and Ch- mice compared to NonTg and ChN, respectively (Fig. 3C). Additionally, a genotype by diet interaction (*F*_(1, 12)_ = 4.966, *p* = 0.0457) revealed that 3xTg-AD Ch- mice gained more weight than 3xTg-AD ChN (*p* = 0.0003) and NonTg Ch- (*p* = 0.0022) mice. The % change in body weight from baseline to the end of study (Fig. 3D) was also higher in 3xTg-AD compared to NonTg groups (*F*_(1, 12)_ = 8.757, *p* = 0.0119) and between Ch- mice and ChN mice (*F*_(1, 12)_ = 13.75, *p* = 0.0030). These results were as expected, as these mice were a subset of those used in [11,12], and further emphasize the relationship of a Ch- diet with weight gain in 3xTg-AD and NonTg mice. Further, we measured plasma choline levels at 7 and 12 months of age. At 7 months (Fig. 3E), 3xTg-AD mice had lower circulating blood choline levels than NonTg mice (*F*_(1, 7)_ = 66.55, *p* < 0.0001). Additionally, a Ch- diet led to lower plasma choline levels compared to mice on the ChN diet (*F*_(1, 7)_ = 9.839, *p* = 0.0165). At 12 months (Fig. 3F), Ch- mice continued to have lower plasma choline levels (*F*_(1, 7)_ = 49.03, *p* = 0.0002). Finally, we performed a GTT to assess glucose metabolism. In healthy individuals, glucose levels spike following glucose consumption and return to baseline levels as insulin shuttles glucose into cells [35]. We found that mice on a Ch- diet had higher glucose levels than those on a ChN diet (*F*_(1, 12)_ = 5.467, *p* = 0.038). A significant timepoint by diet interaction (Fig. 3G; *F*_(7, 84)_ = 2.183, *p* = 0.044) revealed that mice in the Ch- group had higher glucose levels at 15 (*p* = 0.005), 45 (*p* = 0.033), 60 (*p* = 0.043), and 90 (trending; *p* = 0.064) minutes post-glucose injection than those in the ChN group. These findings were reflected in an area under the curve (AUC) analysis, which revealed that glucose AUC was higher in the Ch- mice (Fig. 3H; *F*_(1, 12)_ = 5.736, *p* = 0.0338) compared to the ChN counterparts. Together, these data demonstrate that a Ch- diet contributes to glucose metabolism impairments, consistent with published reports [12,13]. Additionally, the reduction of choline in 3xTg-AD at 7 months, when AD-like pathology is not yet advanced, suggests that early reductions in choline may be a contributor to pathological progression as highlighted below.

**Figure 3.**
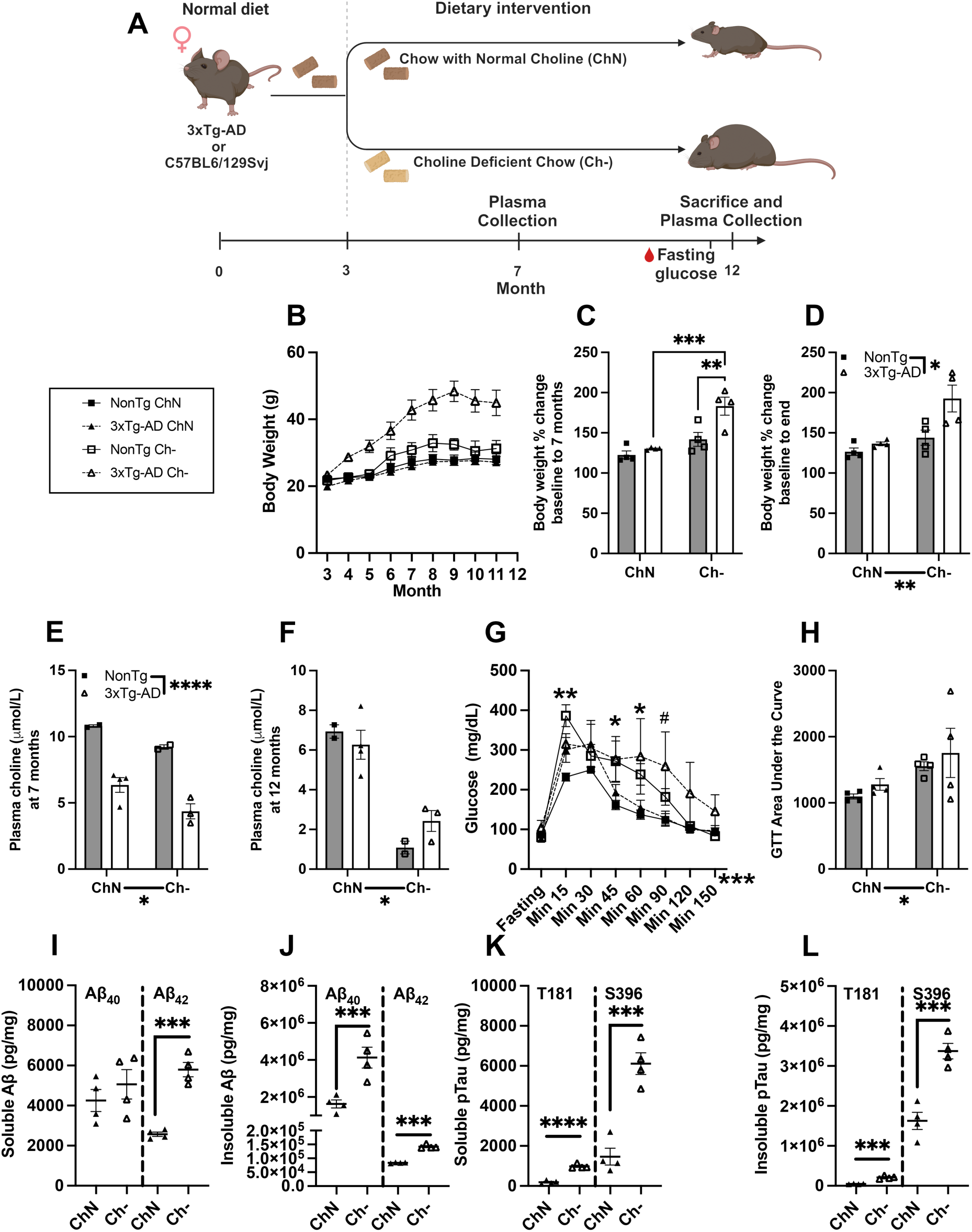
Mice fed a choline deficient diet paralleled obese participants. **(A)** Timeline of dietary intervention. **(B)** Body weight changed across the months of dietary intervention. **(C)** At 7 months, 3xTg-AD Ch- mice gained more weight than NonTg- Ch- and 3xTg-AD ChN. **(D)** At 12 months, Ch- and 3xTg-AD gained more weight than ChN and NonTg, respectively. At 7 **(E)** and 12 **(F)** months, mice on the Ch- diet had lower circulating choline levels than mice on the ChN diet. **(G)** Mice on the Ch- diet had higher glucose levels during the glucose tolerance test (GTT). **(H)** Mice on the Ch- diet had a higher area under the curve (AUC) for the GTT than the ChN counterparts. **(I)** Soluble Aβ40 (left) did not differ based on diet, but soluble Aβ42 (right) was higher in 3xTg-AD mice on a Ch- diet. **(J)** Insoluble Aβ40 (left) and Aβ42 (right), **(K)** soluble pTau T181 (left) and S396 (right), and **(L)** insoluble pTau T181 (left) and S396 (right) were all higher in in 3xTg-AD mice on a Ch- diet. Data are reported as means ± SEM. #P < 0.06, \**p* < 0.05, \*\**p* < 0.01, \*\*\**p*<0.001, \*\*\*\**p* < 0.0001.

### Dietary choline deficiency exacerbates AD pathology in the 3xTg-AD mouse

To confirm the impact of varying dietary choline intake on AD pathological progression, we assessed amyloidosis and hyperphosphorylated tau levels in cortices of the 3xTg-AD mouse. Given that 3xTg-AD mice contain human AD familial mutations, we measured human versions of AD pathology and did not include NonTg in this analysis. For Aβ, we assessed both soluble and insoluble Aβ_40_ and Aβ_42_. There was no significant difference in soluble Aβ_40_ (Fig. 3I Left) between 3xTg-AD mice on the Ch- and ChN diets. However, soluble Aβ_42_ (Fig. 3I Right) levels were significantly higher in mice on a Ch- diet (*t*_(6)_ = 8.565, *p* = 0.0001). Insoluble Aβ_40_ (Fig. 3J Left; *t*_(6)_ = 4.162, *p* = 0.0059) and Aβ_42_ (Fig. 3J Right; *t*_(6)_ = 16.32, *p* < 0.0001) levels were also elevated in 3xTg-AD Ch- mice compared to those on the ChN diet. For markers of pathological tau hyperphosphorylation, we measured soluble and insoluble pTau at two pathological epitopes [11,12,25], T181 and S396. Soluble T181 (Fig. 3K Left; *t*_(6)_ = 11.49, *p* < 0.0001) and S396 (Fig. 3K Right; *t*_(6)_ = 6.040, *p* = 0.0009) and insoluble T181 (Fig. 3L Left; *t*_(6)_ = 8.578, *p* = 0.0001) and S396 (Fig. 3L Right; *t*_(6)_ = 6.785, *p* = 0.0005) were all significantly elevated in 3xTg-AD on the Ch- diet compared to those on the ChN diet. Together, this data validates that AD pathology is elevated following a Ch- diet, highlighting the detrimental brain effects in addition to peripheral metabolic dysfunction.

### Dietary choline deficiency contributes to widespread inflammation

To assess the impact of dietary choline deficiency on inflammation, we examined the same cytokines that were assessed in the obese and healthy participants in a cohort of 3xTg-AD mice that were on Ch- versus ChN diets. We identified several cytokines that were significantly elevated in 3xTg-AD mice and/or by a Ch- diet (Fig. 4A). 3xTg-AD mice had significantly higher levels of IFN-γ (Fig. 4B; *F*_(1,10)_ = 6.715, *p* = 0.0269), IL-1β (Fig. 4C; *F*_(1,10)_ = 13.59, *p* = 0.0042), and IL-2 (Fig. 4D; *F*_(1,10)_ =11.15, *p* = 0.0075). A Ch- diet significantly elevated these cytokine levels regardless of genotype: (IFN-γ (*F*_(1,10)_ = 44.82, *p* < 0.0001); IL-1 β (*F*_(1,10)_ = 33.15, *p* = 0.0002); IL-2 (*F*_(1,10)_ = 14.14, *p* = 0.0037). There were no significant differences in diet or genotype in IL-4 levels (Fig. 4E). 3xTg-AD mice also showed elevated IL-5 (Fig. 4F; *F*_(1,10)_ = 7.541, *p* = 0.0190) and IL-6 (Fig. 4G; *F*_(1,10)_ = 11.00, *p* = 0.0069). IL-5 (*F*_(1,10)_ = 18.40, *p* = 0.0013) and IL-6 were also elevated by a Ch- diet (*F*_(1,10)_ = 17.38, *p* = 0.0016). IL-10 was elevated by the Ch- diet (Fig. 4H; *F*_(1,10)_ = 7.203, *p* = 0.0213). 3xTg-AD mice exhibited elevated IL-12 p70 (Fig. 4I; *F*_(1,10)_ = 18.11, *p* = 0.0014), and IL-12 p70 was elevated by the Ch- diet (*F*_(1,10)_ = 42.95, *p* < 0.0001). The Ch- diet elevated IL-13 (Fig. 4J; *F*_(1,10)_ = 6.934, *p* = 0.0233). Lastly, TNF-α levels were higher in in 3xTg-AD (Fig. 4K; *F*_(1,10)_ = 10.94, *p* = 0.0070) and following the Ch- diet (*F*_(1,10)_ = 32.47, *p* = 0.0001). These findings indicate that a deficiency in dietary choline increases inflammation, which aligns with our observations in obese individuals. Notably, this includes cytokines such as IL-6, IFN-γ, and TNF-α, which are associated with both diabetes and dementia. [11,31,32,36,37].

**Figure 4.**
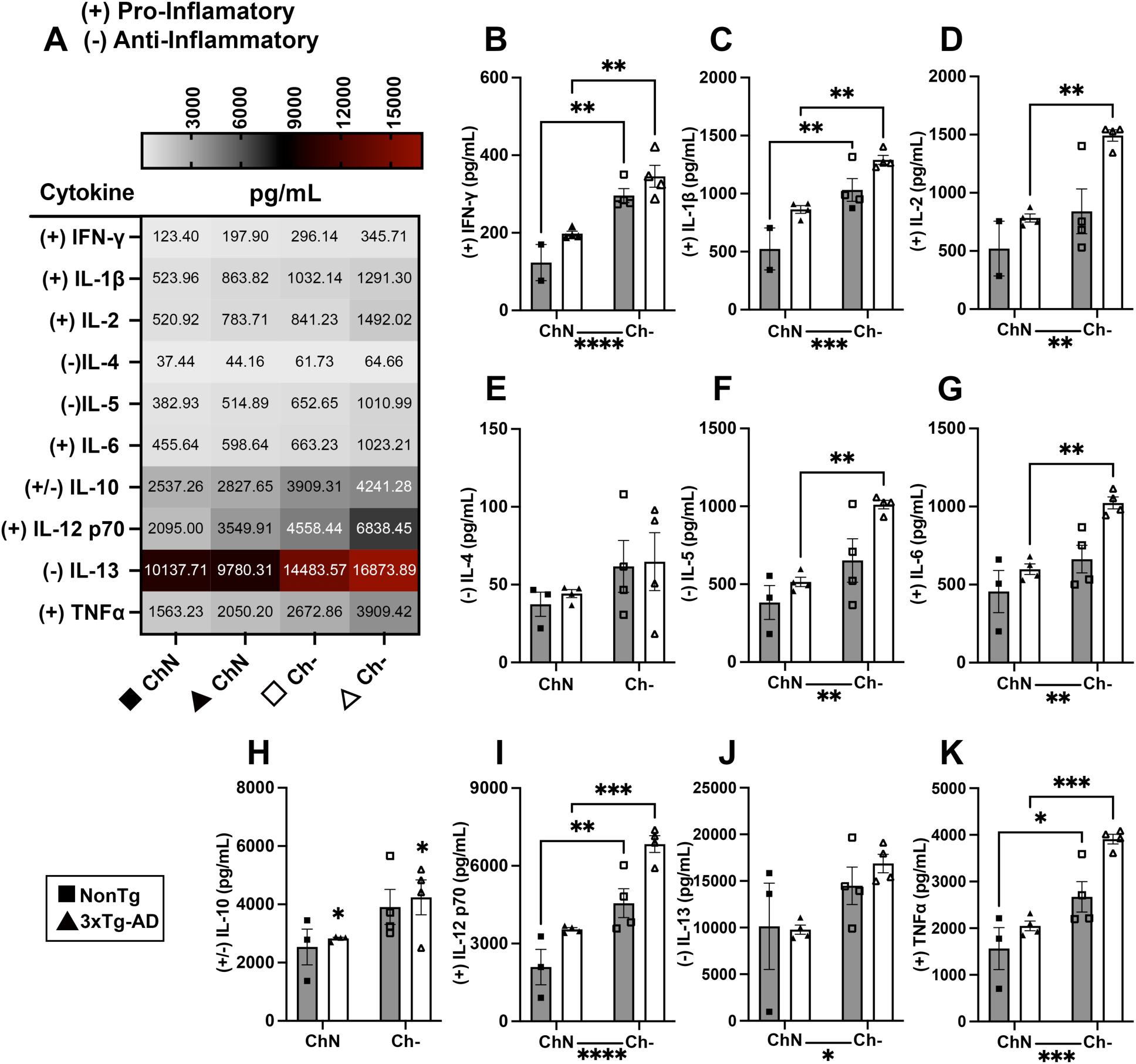
Eleven cytokines were assessed in 3xTg-AD mice on a Ch- diet. **(A)** A heatmap illustrating the changes in the 11 cytokines assessed. **(B)** IFN-γ, **(C)** IL-1β, and **(D)** IL-2 were higher in mice fed a Ch- diet. **(E)** IL-4 didn’t differ based on diet. **(F)** IL-5 and **(G)** IL-6 were higher in mice on the Ch- diet. **(H)** IL-10 was higher in 3xTg-AD mice than in NonTg. **(I)** IL-12 p70, **(J)** IL-13, and **(K)** TNF-α were higher in mice on the Ch- diet. Data are reported as means ± SEM. \**p* < 0.05, \*\**p* < 0.01, \*\*\**p*<0.001, \*\*\*\**p* < 0.0001.

## Discussion

Here, we demonstrate a relationship between early to mid-life obesity, IR, circulating choline levels, and cytokine inflammation profiles, emphasizing their potential as risk factors for cognitive decline and AD. Our results show that choline levels were significantly lower in obese participants compared to those of healthy weight. Importantly, several metabolic indicators were elevated in obese participants, and both body composition markers (BMI and % Body fat) and insulin sensitivity markers (insulin levels and HOMA-IR) showed negative correlations with choline levels. Obese participants also exhibited dysregulated inflammation compared to healthy BMI controls, indicated by the elevation of 11 cytokines. Additionally, obese participants had elevated aldolase B and sorbitol dehydrogenase, liver enzymes involved in sugar metabolism, and these elevations were correlated with reduced choline levels, paralleling our past findings in 3xTg-AD mice on a Ch- diet [12]. A Ch- diet in 3xTg-AD mice resulted in weight gain, impaired glucose metabolism, exacerbated AD pathology, and elevated inflammation. Thus, the findings in obese human participants and 3xTg-AD mice on a Ch- diet show numerous parallels and support the role of low choline in the potential development of metabolic and inflammatory dysfunctions associated with obesity, which may ultimately increase the risk of disorders such as AD.

We identified IR in obese participants and impaired glucose metabolism in Ch- 3xTg-AD mice. Both obese participants and Ch- 3xTg-AD mice exhibited reduced circulating choline levels, emphasizing the potential links between metabolic dysfunction and choline deficiency. IR disrupts glucose metabolism and can lead to pancreatic β-cell dysfunction, hampering the pancreas’ ability to maintain normal glucose levels [4]. IR is prevalent in prediabetes, T2D, and AD, playing a pivotal role in the pathology of each condition by disrupting cellular metabolism and promoting inflammation [38,39]. Individuals with obesity are seven times more likely to develop T2D compared to those with a healthy weight [2]. T2D is diagnosed when fasting glucose exceeds 126 mg/dL, 2-hour post-glucose exceeds 200 mg/dL, or HbA1c is ≥6.5% [40]. Obesity further contributes to IR through increased adiposity, particularly visceral fat, which secretes pro-inflammatory cytokines and fatty acids that disrupt normal metabolic processes [3]. This chronic inflammation induces IR not only in peripheral tissues but also in the brain [2,41]. Historically, the relationship between cognitive impairment and metabolic diseases was underappreciated, but emerging epidemiological evidence now highlights a significant association between these conditions [42,43]. Studies suggest that IR, chronic low-grade inflammation, and disrupted brain insulin signaling may serve as shared pathogenic mechanisms between obesity, T2D, and AD [42–45]. T2D is associated with a 10-year reduction in life expectancy and increased risks of cardiovascular disease, kidney failure, vision loss, and limb amputations [46]. Additionally, it raises the risk of AD by promoting IR in the brain, which disrupts tau phosphorylation and impairs Aβ clearance, contributing to cognitive decline [10]. These findings emphasize the critical importance of early intervention during the obesity stage to prevent progression to T2D and AD. Targeting common pathways—such as insulin signaling, inflammation, and metabolic dysregulation—could offer valuable therapeutic opportunities for both T2D and AD, potentially slowing disease progression and improving the quality of life for individuals affected by these conditions.

Choline deficiency may be intricately linked to obesity, impaired glucose metabolism, and an increased risk of AD, due to its essential roles in regulating various metabolic functions [12,15,16,47]. Reduced choline levels are associated with a higher prevalence of AD [14,19,20]. Choline deficiency is widespread; on average, males consume about 402-mg/day of choline, while females average only 278-mg/day, both of which are significantly below the recommended daily intake of 550- and 425-mg/day, respectively [48,49]. In our study, circulating choline levels were lower in obese individuals compared to those with a healthy BMI, with females exhibiting lower choline levels than males, despite typically having higher endogenous choline production due to estrogen’s activation of the PEMT enzyme [50]. Strong correlations were observed between reduced plasma choline levels and various metrics of obesity and IR, with higher BMI and %Body fat correlating with decreased choline levels. Additionally, lower choline levels were associated with higher insulin and HOMA-IR scores. Notably, our female participants had a higher %Body fat, consistent with previous research suggesting that women may experience more pronounced metabolic and inflammatory effects from obesity due to higher %Body fat [3]. Additionally, decreased choline is associated with higher pathological markers of AD, such as increased neuritic plaque density and a higher Braak stage [11,20]. Reduced choline levels also correlate with worse AD clinical outcomes, including lower Mini-Mental State Examination scores and decreased brain weight [11]. Taken together, these findings suggest that low choline levels in adulthood may contribute to IR, which could, in turn, promote the development of AD later in life. Lastly, the lower choline levels observed in women may contribute to their increased risk of developing AD [7].

Alongside choline deficiency, obesity and IR are characterized by inflammatory components that further complicate metabolic health [3]. Individuals with obesity exhibit elevated levels of both pro-inflammatory and anti-inflammatory cytokines, contributing to a persistent state of chronic inflammation [3,21,51]. This chronic systemic inflammation, triggered by obesity, plays a central role in the development of IR, β-cell dysfunction, and ultimately T2D as well as conditions like AD [2]. Inflammation in adipose tissue drives IR by impairing insulin receptor function or insulin secretion, which alters glucose sensitivity [2]. Moreover, inflammation exacerbates the development of AD pathology through increased apoptosis and reduced synaptic activity [41]. Inflammatory responses disrupt insulin signaling and its downstream pathways [10]. IR in the brain promotes the release of pro-inflammatory cytokines [41]. Our analysis revealed that eleven cytokines were significantly elevated in obese participants compared to healthy controls, with pro-inflammatory cytokines, such as INF-γ, IL-1β, and TNF-α being notably increased. INF-γ induces adipose tissue inflammation in T2D [52], while IL-1β and TNF-α are associated with IR [3]. Similarly, IL-10, while promoting IR in adipocytes, enhances insulin sensitivity in skeletal muscle [3]. IL-2 has also been linked to both IR and elevated blood glucose levels [33]. IFN-γ induces the release of additional inflammatory signals from adipose tissue [52]. IL-8, a pro-inflammatory cytokine, is elevated in T2D patients and correlates with poorer glycometabolic profiles [34]. IL-12p70 is associated with endothelial dysfunction and cardiovascular disease in diabetes [53]. Interestingly, despite the dominance of pro-inflammatory markers, anti-inflammatory cytokines like IL-4, IL-5, and IL-13—which counteract inflammation in adipose tissue and enhance insulin sensitivity—were also elevated in obese participants [3]. This suggests that the body attempts to counterbalance the inflammatory response. However, the imbalance between pro- and anti-inflammatory signals only further exacerbates the chronic inflammatory environment [35]. Notably, elevated levels of several of these cytokines—especially IL-6, IFN-γ, and TNF-α—have been proposed as links between IR and dementia, with higher levels associated with increased AD pathological burden [3].

Choline deficiency worsens obesity-related inflammation, potentially disrupting lipid metabolism and triggering the release of inflammatory mediators [21]. Using the 3xTg-AD animal model and NonTg control mice, we further identified that choline deficiency parallels the inflammatory changes observed in obese participants. Choline deficiency may worsen inflammatory responses and influence the progression of AD pathology, exacerbating amyloid and tau pathogenesis in the 3xTg-AD mouse model. Additionally, another study from our lab demonstrated that choline supplementation in the APP/PS1 mouse model increased circulating choline levels while reducing Aβ and TNF-α levels in the brain [11,18]. Together, these findings add to the growing body of data that adequate choline levels are vital for preventing chronic inflammation and metabolic disorders, emphasizing the complex, multifaceted nature of inflammation in obesity and its role in the progression of metabolic dysfunction. This is supported by other studies in both humans and rodents that suggest higher dietary choline intake reduces the risk of IR [13], T2D [15], and impaired glucose tolerance, and alterations in insulin-degrading proteins that increase the likelihood of IR [12]. However, other evidence points to a more nuanced relationship between choline and IR. While choline levels decrease in non-obese individuals as they transition from insulin sensitivity to IR, higher levels of choline phospholipids are observed during the progression to IR, suggesting that choline metabolism dysregulation may contribute to IR development [16]. Moreover, studies highlighting the increased risk of obesity, IR, and other health issues focus on the choline metabolite trimethylamine N-oxide (TMAO), which is produced by certain gut microbiome species more prevalent in individuals with diets high in red meat and low in fiber [54,55]. While our study indicates that higher choline levels improve metabolic health, and when considered alongside the whole of the literature, choline intake should complement – not replace – a balanced diet.

In our previous work, we identified the liver enzymes aldolase B and sorbitol dehydrogenase as being dysregulated by a Ch- diet [12]. Here, we found that these enzymes are increased in obese participants compared to healthy controls, and that circulating choline levels negatively correlate with rises in these enzymes. These enzymes play a crucial role in sugar metabolism: aldolase B metabolizes fructose, while sorbitol dehydrogenase metabolizes glucose [30]. Elevated levels of both enzymes in the blood can indicate liver dysfunction [56,57]. Furthermore, increased aldolase B is associated with reduced insulin production [58], and raised sorbitol dehydrogenase levels are found in T2D, contributing to oxidative stress and vascular damage associated with the disease [59]. Interestingly, weight loss can reverse the increase in aldolase B [60], suggesting potential restoration of liver health. Given choline’s role in maintaining liver function and metabolic health, the elevation of aldolase B and sorbitol dehydrogenase levels suggest a potential mechanism through which reduced choline contributes to the metabolic dysfunction seen in T2D, which could also increase AD risk.

Collectively, these findings underscore the significant impact of dietary and circulating choline on obesity and associated metabolic disturbances. Notably, we observed a marked overlap between the metabolic and inflammatory profiles of obese participants with both a control and an AD mouse model on a choline deficient diet. This convergence, alongside research linking IR with AD, strengthens the growing evidence that these diseases share overlapping metabolic and inflammatory changes. Both conditions are exacerbated by low choline levels, indicating that adequate choline may serve as a key mechanism for preventing the metabolic dysregulation contributing to the onset and progression of IR, T2D, and AD. Additionally, considering sex differences in metabolic health is crucial for tailoring effective interventions. Addressing choline deficiencies early could help mitigate or even prevent these disorders, which are increasingly prevalent.

## Abbreviations

AD: Alzheimer’s Disease
Aβ: amyloid beta
NFT: neurofibrillary tau tangles
BMI: body mass index
U.S.: United States
PEMT: phosphatidylethanolamine N-methyltransferase
T2D: type 2 diabetes
IR: insulin resistance (IR)
IL: Interleukin
TNF: Tumor necrosis factor
BIA: bioelectrical impedance analysis
APP: amyloid precursor protein
PS1: presenilin 1
NonTg: non-transgenic
ChN: diet with adequate choline
Ch-: choline-deficient diet
GTT: glucose tolerance test
pTau: phosphorylated Tau
HOMA-IR: Homeostatic Model Assessment for IR
AUC: area under the curve
IFN: interferon
M: Mean
SEM: Standard error mean

## Credit authorship contribution section

**Wendy Winslow:** Writing – original draft, ELISAs, Conceptualization. **Jessica Judd:** Writing – original draft, Animal experiments, Analysis, Conceptualization. **Savannah Tallino:** Animal experiments, writing and editing the manuscript. **Lori R. Roust:** IRB, Human data collection. **Eleanna De Filippis:** IRB, Human data collection**. Brooke Brown:** IRB, Human data collection. **Christos Katsanos:** Human data analysis, Writing and editing the manuscript. **Ramon Velazquez:** Writing – review & editing, Supervision, Investigation, Funding acquisition, Conceptualization.

## Ethics approval statement

The human studies were approved by the Institutional Review Board at Mayo Clinic (IRB #: 20-003294). All experimental procedures on animals were approved by Arizona State University IACUC and carried out according to the approved guidelines.

## Acknowledgements

Fig. 2 partly Created in BioRender.

## Data availability

Data will be made available upon reasonable request.

